# Synthetic cell-cycle regulation identifies Mif2^CENP-C^ as a CDK phospho-target at the kinetochore

**DOI:** 10.1101/2023.03.24.534130

**Authors:** Cinzia Klemm, Guðjón Ólafsson, Peter H. Thorpe

## Abstract

Protein phosphorylation regulates multiple cellular processes including cell-cycle progression, which is driven by highly conserved cyclin-dependent kinases (CDKs). CDKs are controlled by the oscillating levels of activating cyclins and the activity peaks during mitosis to promote chromosome segregation. However, with some exceptions, we do not understand how the multitude of CDK-phosphorylated residues within the proteome drive cell-cycle progression nor which CDK phosphorylation events are necessary. To identify yeast proteins whose phospho-regulation is most critical for cell-cycle progression, we created a synthetic CDK complex and systematically recruited this to proteins involved in chromosome segregation using the Synthetic Physical Interactions (SPI) method. We found that targeted recruitment of synthetic CDK to the centromeric protein Mif2^CENP-C^ leads to enrichment of Mif2^CENP-C^ at centromeres and arrested cells in late mitosis. We then identified putative CDK consensus sites on Mif2^CENP-C^ which aid Mif2^CENP-C^ localisation at centromeres and showed that CDK-dependent Mif2^CENP-C^ phosphorylation is important for its stable kinetochore localisation.

**Summary:** To identify cellular sites of functional cell cycle phospho-regulation we generated a synthetic cyclin-dependent kinase which can be recruited to any given GFP-tagged protein. Using this system with a set of proteins involved in chromosome segregation, we identified Mif2^CENP-C^ as a kinetochore target of CDK and show that CDK stabilises Mif2’s kinetochore localisation.

## Introduction

CDK-mediated protein phosphorylation is the key mechanism underlying cell-cycle progression (Alberghina et al., 2012). Different cyclins recruit the kinase to its substrate proteins, dependent on the cell-cycle stage (De Bondt et al., 1993; Malumbres, 2014). A single kinase, Cdc28^CDK1^, forms the active part of the cell cycle-promoting CDK complex in the budding yeast *S. cerevisiae,* which can be functionally complemented with its human counterpart CDK1 (Lee & Nurse, 1987; Hamza et al. 2015). Coupled with the genetic tools available in yeast, this system makes a highly conserved, yet tractable model organism to study CDK phospho-regulation systematically. CDK activity is tightly regulated by cyclin-binding and peaks in early mitosis, when Cdc28^CDK1^ is bound to M-phase cyclins or Clb^CyclinB^ proteins, to promote progression from metaphase into anaphase and chromosome segregation (Chee & Haase, 2010; Loog & Morgan, 2005; Rahal & Amon, 2008). During anaphase, Clb^CyclinB^ is targeted for degradation by the anaphase-promoting complex (APC), to abolish CDK activity (Herzog et al., 2009; Lim et al., 1998). When CDK activity falls at the end of mitosis, CDK-mediated phosphorylation is reversed by cell cycle phosphatases, and in budding yeast, the Cdc14 phosphatase dephosphorylates the majority of CDK substrates (Kuilman et al., 2015; Stegmeier & Amon, 2004).

CDK activity is strictly regulated by cyclin association, as cyclin-binding promotes conformational changes within the active center of the kinase subunit Cdc28 (Jeffrey et al., 1995). In *S. cerevisiae*, nine cyclin subunits activate Cdc28 at distinct times during the cell cycle, which can be subdivided into G1 cyclins (Cln1-4), S/G2 cyclins (Cln5-6) and mitotic cyclins (Clb1-3). None of the cyclins are essential on their own to promote cell-cycle progression, however, deleting all cyclins important for progression into the next phase halts the cell cycle. Previous research which aimed to abolish cyclin requirement led to the design of a G1 cyclin-bypassing mutant of Cdc28 (Levine et al., 1999), however S/G2/M phase cyclins remained indispensable. In the fission yeast *S. pombe*, a synthetic, minimal cell cycle can be driven by a single cyclin-CDK fusion protein, consisting of the mitotic cyclin Cdc13 (Clb2 in *S. cerevisiae*) fused to the N-terminus of the kinase subunit (Coudreuse & Nurse, 2010). In *S. cerevisiae* G1 cyclin-specificity is essential for morphogenetic development, as replacing G1 cyclins with mitotic cyclins prevents budding (Pirincci Ercan et al., 2021).

One target complex of CDK phosphorylation is the kinetochore, a megadalton protein complex consisting of more than 60 subunits (Biggins, 2013; Kitazono & Kron, 2002). The kinetochore attaches chromosomes to the mitotic spindle and can be divided into the constitutive centromere associate network (CCAN), which binds to the centromeric region of chromosomes, and the KMN network which attaches to spindle microtubules. Phospho- regulation is crucial for kinetochore function and indispensable for correct chromosome segregation (Klemm et al., 2021). Many kinetochore proteins contain CDK consensus sites, however, only a few phosphorylation events are functionally characterised (Bock et al., 2012; Böhm et al., 2021; Gutierrez et al., 2020; Li & Elledge, 2003). Besides CDK, other mitotic kinases such as Ipl1^AuroraB^, Mps1, DDK and Cdc5^Polo^ target kinetochore subunits (Hinshaw et al., 2017; Keating et al., 2009; Maure et al., 2007; Mishra et al., 2019; Mishra & Basrai, 2019; Ólafsson & Thorpe, 2020; Shimogawa et al., 2006), indicating a complex interplay of phospho- regulators important for kinetochore assembly and function.

A highly phospho-regulated component of the budding yeast CCAN is the conserved kinetochore protein Mif2^CENP-C^ (Hinshaw et al., 2023; Hinshaw & Harrison, 2019; Peng et al., 2011). Mif2^CENP-C^ forms a homodimer via its C-terminal Cupin dimerization domain (Cohen et al., 2008). A central domain of Mif2^CENP-C^ interacts with the centromeric histone Cse4^CENP-A^, whereas its N-terminal domain binds to Mtw1^MIS12^ of the KMN network (Cohen, 2008; Dimitrova et al., 2016). The interaction between Mif2^CENP-C^ and Cse4^CENP-A^ is positively regulated by the tension-sensing kinase Ipl1, and casein kinase CK2 (Peng, 2011), whereas Mtw1-binding is regulated by Mif2^CENP-C^ autoinhibition (Killinger et al., 2020). Mif2^CENP-C^ further contains a PEST domain, enriched with the amino acids proline, glutamate, serine and threonine, which is commonly associated with a shorter half-life of proteins as it serves as a proteolytic signal (Rechsteiner & Rogers, 1996; Scott et al., 1986). In human cells, the PEST sequence is involved in CENP-C binding to CENP-HIKM, and CENP-HIK are human homologs of the budding yeast CTF3 complex proteins Mcm16, Ctf3 and Mcm22 (Klare et al., 2015). Mif2^CENP-C^ phosphorylation by Ipl1^AuroraB^ and CK2 impacts protein stability (Peng, 2011). Human CENP-C has been identified as a target of CDK, and the PEST domain of *S. cerevisiae* is phosphorylated by Cdc5^Polo^ and Dbf2, mitotic kinases which often recognise CDK-phosphorylated proteins (Hinshaw, 2023). However, there is no direct experimental evidence of CDK-mediated Mif2^CENP-C^ phosphorylation in budding yeast.

To identify if Mif2^CENP-C^ is a target of CDK phosphorylation, we designed an ectopically expressed, synthetic CDK complex with a strong binding affinity to GFP-tagged proteins. We tested the activity and specificity of this synthetic kinase by forced protein recruitment to select GFP-tagged proteins involved in chromosome segregation. We found that constitutive recruitment of a mitotic CDK complex to Mif2^CENP-C^ strongly increases its kinetochore localisation and prevents cytokinesis. We further identified Mif2^CENP-C^ as a direct target of CDK and show that Mif2^CENP-C^ phosphorylation supports kinetochore stability.

## Results

Large-scale phospho-proteomic studies have been used to systematically identify global phosphorylation landscapes throughout the cell cycle (Holt et al., 2009; Lanz et al., 2021; Ubersax et al., 2003; Touati et al., 2019). For CDK alone, more than 1000 phospho-sites within over 300 substrate proteins have been reported, underlining its power to drive a multitude of cell cycle processes. Despite the major advances in substrate identification mediated by phospho-proteomics, functional aspects of specific phosphorylation events often remain unclear. Classical approaches to understanding the functional role of CDK phospho- regulation, such as using conditional mutants or deletion of activating cyclins, have identified more general consequences of reduced CDK activity (Hartwell & Smith, 1985; Kuczera et al., 2010; Schneider et al., 1996; Schwob & Nasmyth, 1993). However, these studies seldomly report the effects of phosphorylation of certain complexes or proteins. On the other hand, targeted mutation of CDK consensus motifs is a powerful tool to focus on the impact of specific protein phosphorylation events but requires a good knowledge about CDK-target sites and cannot be easily scaled up to medium or high-throughput studies (Akiyoshi & Biggins, 2010; Böhm et al., 2021; Gutierrez, 2020; Zegerman & Diffley, 2007). We have previously shown that systematic, forced kinase recruitment serves as a powerful tool to unravel functional aspects of protein phosphorylation on a proteome-wide scale (Ólafsson & Thorpe, 2015, 2016; Ólafsson, 2020). To forcibly promote these kinase-substrate interactions, we use the Synthetic Physical Interactions method or SPI. In brief, SPI utilises an ectopically expressed phospho-regulator fused to a small GFP binding protein (GBP or nanobody, (Fridy et al., 2014; Rothbauer et al., 2006) (Figure 1A)). This construct is expressed in strains containing different potential substrate proteins fused with GFP (Huh et al., 2003). As the read-out of SPI is colony growth, with small or absent colonies indicating cell cycle perturbation, we conclude that growth defects highlight crucial substrates which are dynamically phosphorylated and dephosphorylated throughout cell cycle progression.

**Figure 1:**
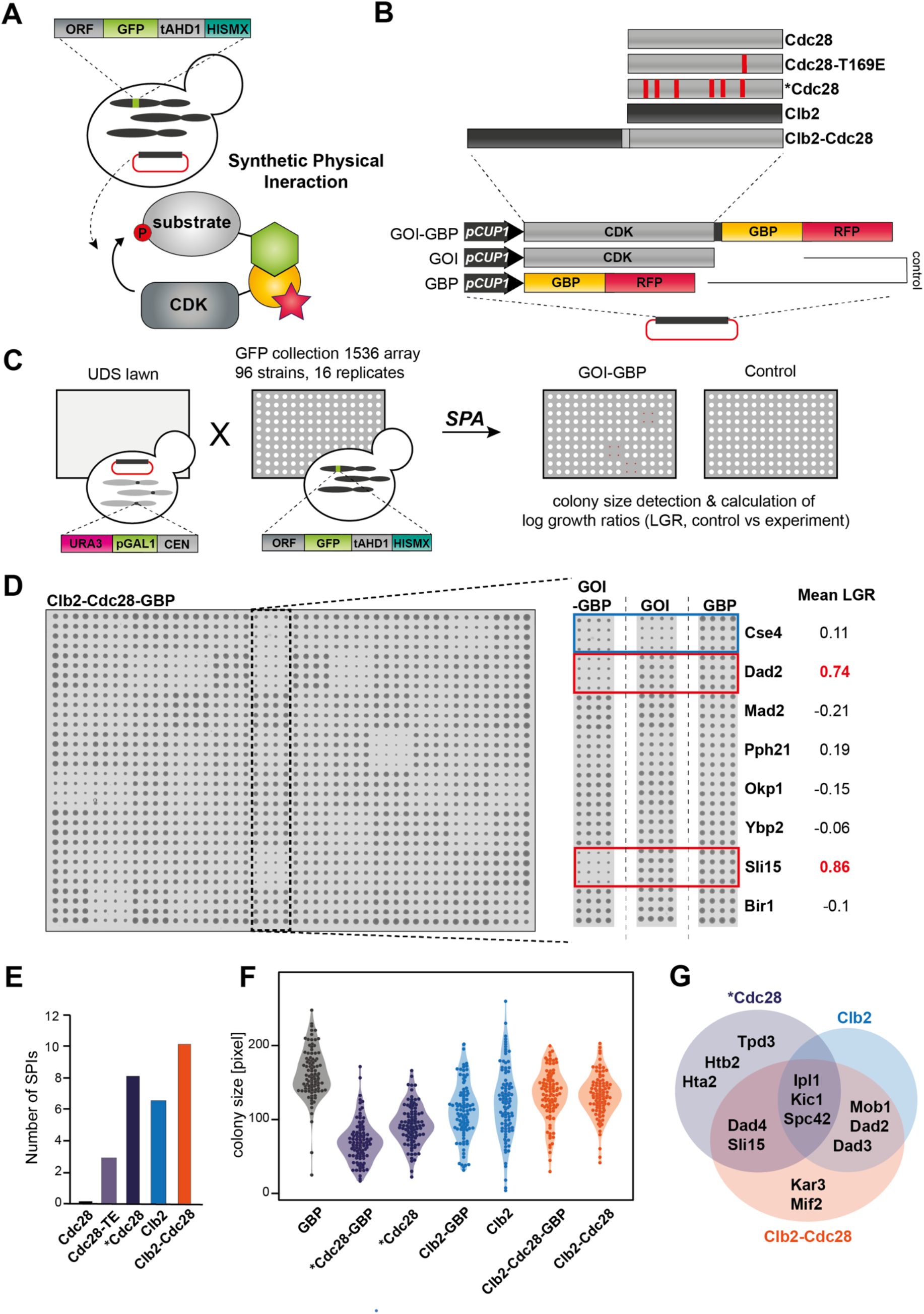
Design of a synthetic CDK suitable for SPI screening. (A) Schematic showing the principle of forced protein interactions using SPI. Due to the strong binding affinity of GBP to GFP, CDK constructs are recruited to query proteins and can phosphorylate substrates. (B) Schematic of CDK constructs encoded by SPI plasmids. All constructs are expressed under a constitutively active CUP1 promoter. SPI plasmids encoded phospho-regulators fused to GBP-RFP. A catalytically dead version of the SPI construct, and plasmids expressing the phospho-regulator or GBP-RFP alone serve as controls. (C) SPI plasmids are transformed into the universal donor strain (UDS) and mated with 89 selected strains of the GFP collection. Diploids are reverted to haploids encoding SPI plasmids and GFP-loci using selective ploidy ablation. (D) Example of SPI screening plate. GFP strains are arranged in 4 replicates and 1636 strains per plate and the effect of phospho-regulator recruitment is assessed by colony size compared to control plates. LGR: Log growth ratio. (E) The number of SPIs for each construct is represented in the bar chart. (F) Raw colony sizes for each screen are represented as boxplots to assess general effects of SPI plasmid expression in GFP strains (1536 colonies per condition). (G) Venn diagram showing the overlap of SPIs between the G1 cyclin-bypassing mutant, Clb2- Cdc28 and Clb2.

In previous studies, we have shown that several kinetochore protein-GBP fusions lead to growth defects in Cdc28-GFP strains, indicating a functional role of CDK phospho-regulation at yeast kinetochores (Ólafsson & Thorpe, 2020). However, Cdc28 is an essential protein and recruiting Cdc28 to kinetochores could prevent phosphorylation of distant (e.g. cytosolic) targets. Furthermore, CDK is strictly regulated and only active when cyclins are bound to Cdc28. Therefore, we wanted to reverse our SPI system and recruit an ectopically expressed, synthetic CDK-GBP fusion to GFP-tagged proteins. We designed and screened different CDK- GBP variants for their ability to be used for SPI (Figure 1B): (1) wildtype Cdc28; (2) Cdc28- T169E, a mutant which promotes phosphorylation of Cdc28 by the CDK activating kinase 1 (Cak1) to facilitate cyclin-binding (Cross and Levine, 1998), (3) a Cdc28 mutant which completely bypasses G1-phase cyclins where Cdc28 acts as an active kinase independent of cyclin binding (Levine et al., 1999), (4) the mitotic cyclin Clb2 and (5) a fusion protein of the M-phase cyclin Clb2 and Cdc28 (Clb2-Cdc28), similar to cyclin-Cdk fusion used in *S. pombe* previously (Coudreuse & Nurse, 2010). All variants were fused to GBP-RFP, to promote association with GFP-tagged proteins and expressed under a constitutively active copper promoter (pCUP1) from low-copy CEN plasmids. We also engineered control plasmids expressing the synthetic CDK without GBP-RFP, which served as controls for ectopic synthetic CDK expression and used a plasmid expressing GBP-RFP on its own as a control for proteins sensitive to forced protein recruitment. Using selective ploidy ablation (SPA, Figure 1C, Reid et al., 2011) we transformed SPI plasmids into 88 strains with different GFP-tagged proteins (Supplementary file 1). These proteins are important for chromosome segregation, some of which are known targets of CDK phospho-regulation. SPI screens were performed in arrays of 1536 colonies on each plate accounting for 16 replicates for each GFP-tagged strain (Figure 1C). Colony sizes were then quantified using the colony measurement tool of *ScreenMill* (Dittmar et al., 2010) and colony sizes of experimental plates, where phospho-regulators are recruited to GFP-tagged proteins, were then compared to GBP and gene-only controls to calculate mean log growth ratios (LGRs) using the ScreenGarden software tool (Klemm et al., 2022) (Figure 1D). Mean LGRs close to zero indicate no significant growth defect based on CDK recruitment compared with controls, whereas mean LGRs > 0.4 account for colonies that grow significantly worse compared to controls; these we term SPIs. We confirmed successful GBP-mediated protein recruitment by colocalisation microscopy (Figure S1A). Strikingly, Cdc28 recruitment did not produce any growth defects when recruited to any of the 88 GFP strains. Cdc28-T169E produced only one SPI with Spc42, a component of the spindle pole body (SPB). The mutant of Cdc28 that bypasses G1 cyclins produced 8 SPIs, Clb2 recruitment led to growth defects with 6 GFP strains and the synthetic CDK (Clb2-Cdc28) produced 10 SPIs (Figure 1E). The majority of SPIs produced by the CDK constructs overlapped between variants and included several kinetochore and SPB proteins (Figure 1F). We further wanted to know whether the expression of variants caused general growth disadvantages in our experimental setup. To this end, we compared colony sizes from experimental GOI control plates to colony sizes of the generally healthy GBP control (Figure 1G). Colony sizes were significantly smaller after recruitment of the G1 cyclin-bypassing mutant, which we confirmed by visually comparing plates. Hence, we chose Clb2-Cdc28 (hereafter referred to as synthetic CDK or sCDK) as the most suitable construct for SPI screening, as we observed the most growth defects when recruited to GFP-tagged proteins without affecting cell-cycle progression and colony size of cells expressing the sCDK GOI control.

Next, we wanted to confirm kinase activity of our sCDK. We designed an additional kinase-dead (kd) version of the sCDK, which is unable to bind ATP (*clb2-cdc28-K40L*) and transformed our plasmids into temperature-sensitive (ts) *cdc28-1* and *clb1,3,4Δ clb2-ts* yeast. We assessed kinase activity by growth in different dilutions at restrictive or permissive temperatures (Figure S2) and found that sCDK and sCDK-GBP were able to rescue Clb2 but not Cdc28 deficiency, as reported in similar studies (Pirincci Ercan et al., 2021). Interestingly, kinase-dead sCDK-GBP was able to partially rescue Clb2 temperature-sensitivity, indicating that clb2- cdc28-kd may bind endogenous Cdc28 to some extent.

We repeated SPI recruitment with both kinase active and dead sCDK constructs and highlighted kinetochore components sensitive to CDK recruitment (Figure 2A and Supplementary data 2). We found that Mif2, Ndc10, Dad2, Dad3, Dad4, the motor proteins Kar3 and Kip3 and the CPC complex (Ipl1 and Sli15) were sensitive to sCDK recruitment. Interestingly, Ame1 and the well-described CDK substrate, Dsn1, were not sensitive to the recruitment of sCDK but led to growth defects after forced association of the kinase-dead complex. Internally tagged Cse4 was sensitive to both, recruitment of inactive and active kinase, whereas a C-terminally tagged version was not. This is consistent with reports that the function of the C-terminally tagged Cse4 has altered function (Wisniewski et al., 2014). Next, we wanted to know if growth defects are mediated by activation of the spindle assembly checkpoint (SAC). The spindle assembly checkpoint, mediated by the Mps1 kinase, prevents chromosome segregation of improperly attached sister chromosomes, which could lead to aneuploidy (Musacchio and Salmon, 2007). Although the SAC is an essential signalling pathway in human cells, the budding yeast SAC is dispensable. We prohibited SAC signalling by deleting the gene encoding the central SAC component Mad3 and repeated SPI recruitment of sCDK (Supplementary file 2). Notably, SPIs with the DAM1/DASH were, at least partially, rescued by *MAD3* deletion, indicating that sCDK recruitment to this complex, triggers SAC activation (Figure 2B). Strikingly, abolishing SAC signalling strongly increased growth defects when sCDK was recruited to Mif2 and promoted sensitivity to kinase-dead CDK recruitment (Figure 2B,C), indicating that constitutive sCDK interacting with Mif2 leads to cell cycle defects independent of the SAC.

**Figure 2:**
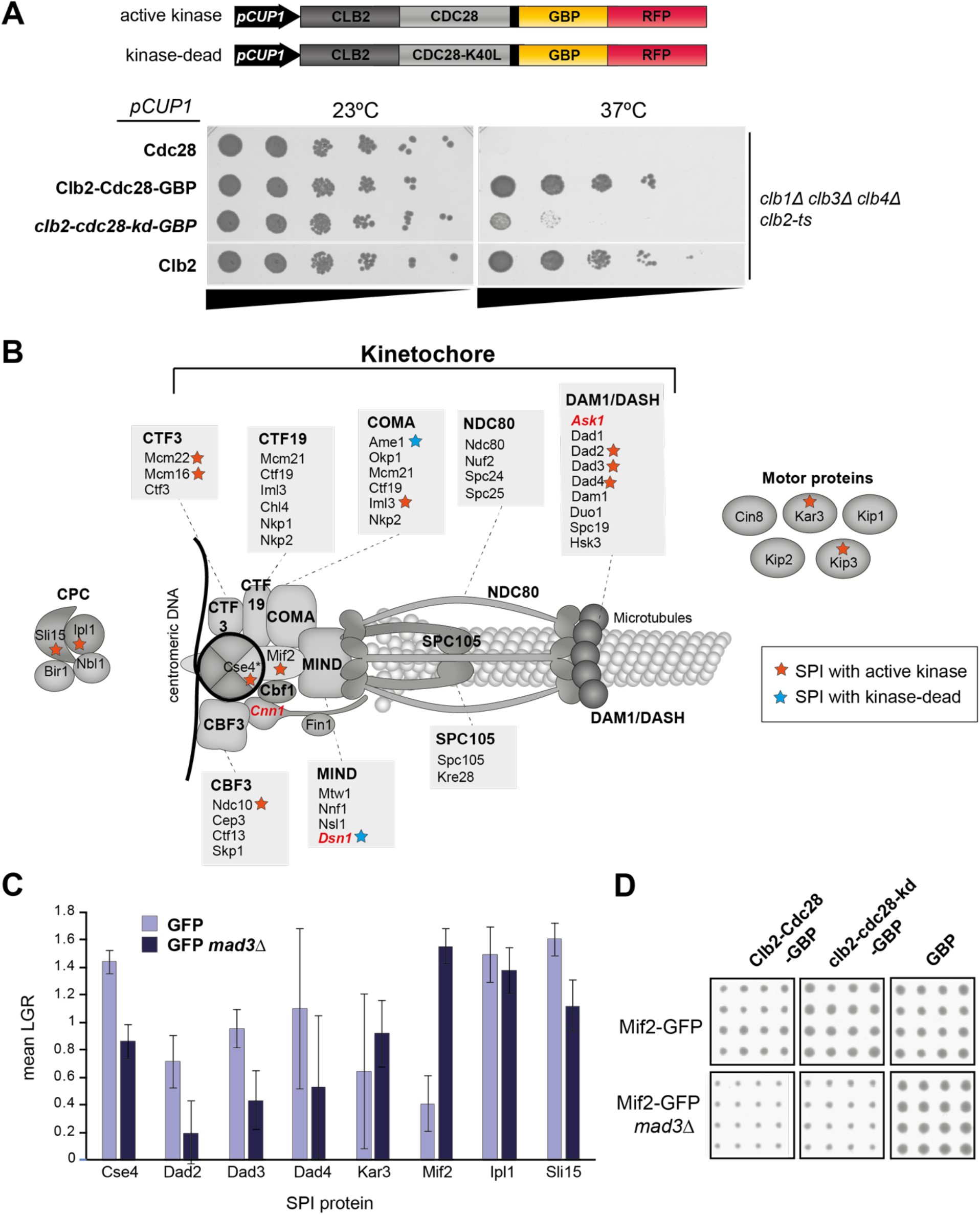
SAC-dependency of kinetochore sCDK SPIs varies between subunits. (A) Active and inactive versions of sCDK were transformed into yeast lacking *CLB1, CLB3* and *CLB4* and encoding a temperature-sensitive (ts) allele of *CLB2* and spot test analysis was performed to assess the capability of SPI constructs to rescue temperature-sensitivity. A *CLB2*- encoding plasmid served as the positive control, whereas Cdc28 overexpression was unable to rescue mitotic cyclin deficiency. Black bars highlight cell dilution (wide end = high number of cells, narrow end = low number of cells). Each step accounts for a ten-fold dilution. (B) CDK SPIs were mapped onto the kinetochore and associated proteins. Orange stars represent SPIs with active CDK, blue stars highlight growth defects with kinase-dead CDK. Known CDK targets are highlighted in red and italics. Cse4* stands for the internally tagged version of Cse4, all other proteins were C-terminally tagged to GFP. (C) SPI screens were repeated with proteins expressing (lilac) or lacking (navy) the *MAD3* gene, encoding a central component of the SAC and LGRs were plotted as a bar chart. Error bars show the standard deviation over 16 replicates. (D) Crops showing colony sizes of Mif2-GFP yeast after recruitment of active or kinase-dead CDK or the GBP-only control in wildtype or *MAD3* deletion backgrounds.

To characterise the immediate cellular response triggered by CDK recruitment to Mif2^CENP-C^, we used CDK-GBP-RFP and GBP-RFP control SPI constructs under the control of the inducible *MET3* promoter (Howell et al., 2020). This system promotes the expression and recruitment of sCDK or GBP to Mif2^CENP-C^ in low methionine conditions but blocks SPI plasmid expression in methionine-rich media. Fluorescence microscopy revealed that, in cells where sCDK was recruited to Mif2 (hereafter referred to as sCDK- Mif2^CENP-C^ cells), Mif2^CENP-C^ fluorescence levels were highly increased compared to GBP recruitment or cells in *pMET3* inhibited conditions (Figure 3A,B and Supplementary file 3). As many sCDK- Mif2^CENP-C^ cells were large-budded cells in late mitosis, we next wanted to understand if sCDK recruitment leads to defective metaphase to anaphase transition. To this end, we used the microtubule-destabilising agent nocodazole to arrest cells before metaphase, generating a synchronised population of cells at the metaphase/anaphase boundary. Cells arrested in nocodazole present a large-budded phenotype; we next quantified the number of large-budded cells to monitor cells’ progression through and exit from mitosis after release from the nocodazole block. We observed that cells expressing sCDK- Mif2^CENP-C^ mostly remained as large-budded cells, whereas control cells (expressing GBP and pMET3) were able to progress and divide (Figure 3C,D and Supplementary data 4). Furthermore, we noted that sCDK- Mif2^CENP-C^ cells quickly progressed from metaphase to anaphase, judged by the interkinetochore distance, but failed to divide into mother and daughter cells (Figure 3C,E and Supplementary file 4). Interestingly, fluorescent tagging of the gamma-tubulin Tub1 revealed that sCDK- Mif2^CENP-C^ cells were not inhibited from spindle disassembly, indicating that mitotic exit is not affected by CDK recruitment as observed by Howell and colleagues (Figure S3D). Moreover, around 6% of large-budded sCDK- Mif2^CENP-C^ cells presented more than two Mif2^CENP-C^ foci, which, judging by the ability of Tub1 association, can assemble associated kinetochore proteins (Figure S3B). Around 30% of sCDK- Mif2^CENP-C^ cells contained weak Mif2^CENP-C^ foci along the mitotic spindle, possibly indicating the presence of lagging chromosomes (Figure S3).

**Figure 3:**
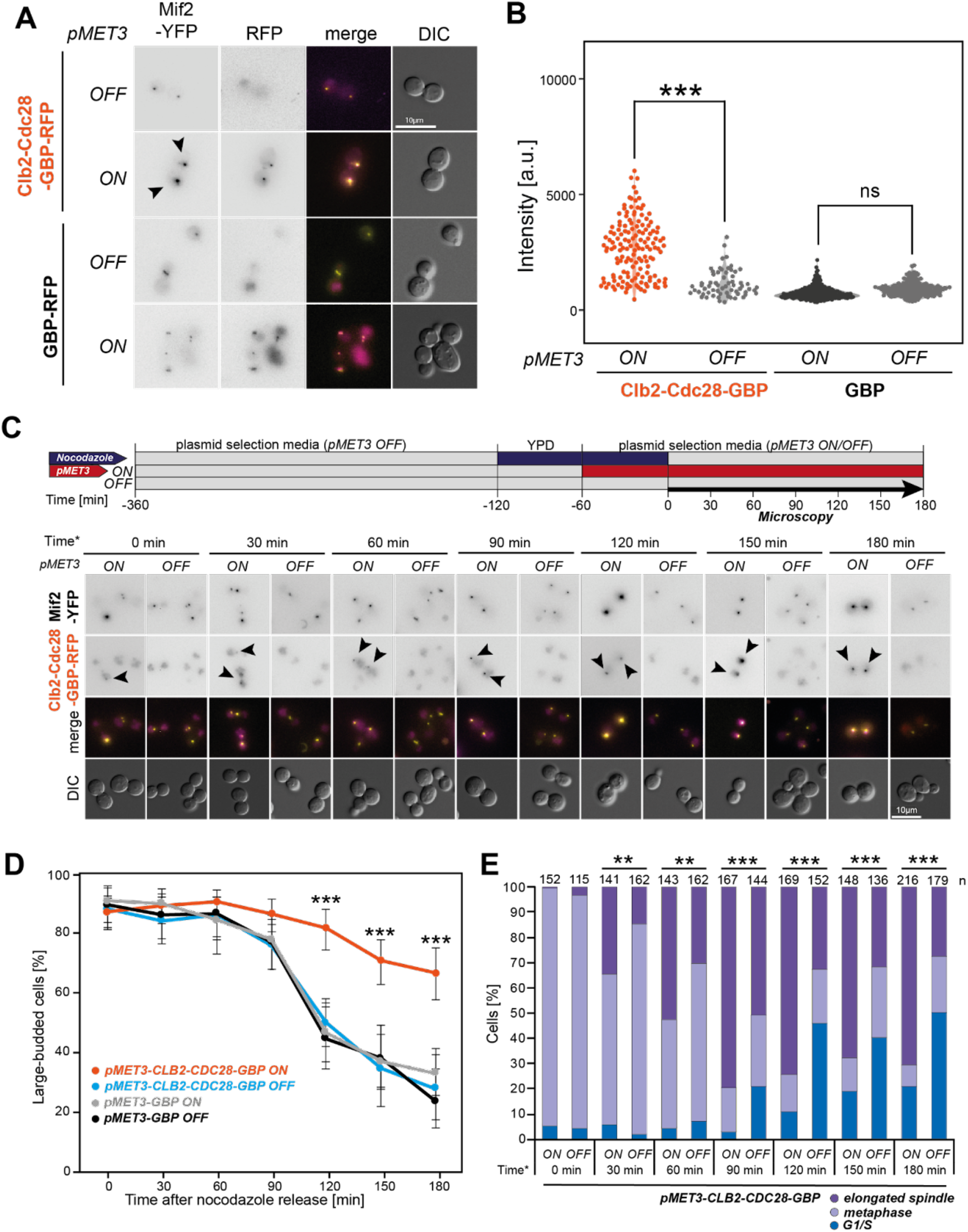
sCDK recruitment to Mif2 leads to kinetochore enrichment and impacts mitotic progression. (A) Micrographs showing conditional protein recruitment of Clb2-Cdc28 to Mif2^CENP-C^. Black arrows highlight enrichment of Mif2^CENP-C^ at kinetochores upon Clb2-Cdc28 recruitment. The scale bar is 10 µm. (B) Quantification of fluorescence intensities of Mif2^CENP-C^ in control conditions (pMET3 OFF or GBP recruitment) and upon Clb2-Cdc28 recruitment. Mif2^CENP-C^ intensities were quantified using the FociQuant software (Ledesma-Fernández & Thorpe, 2015). Statistical significance was assessed using ANOVA (ns = not significant, *** = p-value < 0.005). (C) Schematic and micrographs showing Clb2-Cdc28 recruitment in metaphase cells and mitotic progression. Cells were arrested in metaphase using nocodazole (initial treatment in YPD rather than selection media to promote efficient mitotic arrest). After 1 hour of nocodazole treatment, pMET3 expression was induced in low methionine conditions or repressed at high methionine concentration. Nocodazole was washed out after 2 hours. Cells were then imaged every 30 minutes for 3 hours. The scale bar is 10 µm. (D) Quantification of large-budded cells after nocodazole release in F. Control cells mostly recovered from the large budded phenotype within 2 hours, whereas Clb2-Cdc28 recruitment to Mif2^CENP-C^ arrested cells as large budded. Error bars represent 95% confidence intervals and statistical significance was assessed using Fisher’s Exact test (*** = p-value < 0.005). (E) Quantification of cell cycle stages based on cell shape and kinetochore signals. Clb2-Cdc28 recruitment to Mif2^CENP-C^ led to fast metaphase to anaphase progression, but arrested cells in late mitosis compared to control conditions. Statistical significance was assessed using Fisher’s Exact test (*** = p-value < 0.005).

Our data suggest that CDK phosphorylation of Mif2 aids in the centromeric enrichment of Mif2. Furthermore, we observed that continuous sCDK recruitment to Mif2^CENP-C^ in late anaphase prevents cytokinesis, indicating that dephosphorylation of centromeric CDK targets is crucial for mitotic exit.

To identify whether Mif2^CENP-C^ is indeed a direct target of CDK, we monitored the phosphorylation status of Mif2^CENP-C^ in cells expressing wildtype or temperature-sensitive (ts) alleles of Cdc28 (*cdc28—13* and *cdc28-4*) and the major CDK counteracting phosphatase Cdc14 (*cdc14-1* (Simchen, 1975; Hartwell, 1985; Lörincz A & Reed L, 1986). For Cdc28-ts strains at the restrictive temperature, we observed less retardation of Mif2^CENP-C^ in SDS-PAGE gels containing Phostag agarose, indicating that Mif2^CENP-C^ phosphorylation is decreased when Cdc28 activity is abolished (Figure S4A,B and Supplementary file 5). No change in phosphorylation was observed in either wildtype or *cdc14-1* cells. To see whether this change in phosphorylation directly correlates with Mif2^CENP-C^ levels at kinetochores, we YFP-tagged Mif2^CENP-C^ directly in *cdc28-13* and *cdc28-4* and *cdc14-1* cells. Indeed, we observed reduced Mif2^CENP-C^ kinetochore levels at restrictive temperature in both *cdc28-ts* and *cdc14-ts* strains (Figure S4C,D and Supplementary file 5). Furthermore, an increased number of *cdc28-ts* cells completely lost Mif2^CENP-C^ signals at kinetochores at restrictive temperature (Figure S4E). We note that this phenotype could be explained by the lack of necessity of assembled kinetochores during prolonged G1 arrest. However, Mif2^CENP-C^ is a member of the CCAN and usually remains at centromeres throughout the cell cycle in yeast and human cells (Przewloka et al., 2011; Yan et al., 2019). Furthermore, previous research showed that human CENP-C interacts with CENP-A in a CDK-dependent manner (Watanabe et al., 2019). Given our results, we propose a direct role of CDK-mediated phospho-regulation at Mif2^CENP-C^.

We identified six putative CDK motifs (S/TP or S/TPxR/K) encoded in Mif2^CENP-C^, which are either predicted to be CDK phospho-sites (T345) or were detected as phosphorylated residues by phospho-proteomics (Breitkreutz et al., 2010; Holt et al., 2009; Lanz et al., 2021; MacGilvray et al., 2020; Swaney et al., 2013; Westermann et al., 2003) (Figure 4A). These putative CDK phosphorylation sites are localised in the N-terminal portion of Mif2^CENP-C^, between the N-terminal Mtw1-binding region and the PEST domain (S86) or directly within the PEST domain (S108, T134, S160, T166). The PEST domain also contains a PxL motif, which potentially serves as a recognition site for early and effective dephosphorylation by Cdc14 (Kataria et al., 2018). In addition, a CDK consensus site (TPTR) is present adjacent to a putative AT DNA-binding sequence (T345) (Brown, 1995). We compared these sites to Mif2^CENP-C^ of four homologs in divergent yeast species to assess their conservation (Figure 4B). Notably, T166 is either fully conserved or altered to serine in the four yeast species, and instead of an upstream SP site (S160), negatively charged amino acids (G,D) are present, which could function in a similar way to a phosphorylated residue. In addition, the adjacent PxL motif is also evolutionary conserved in most species. Although the *S. pombe* homolog cnp3 has not been identified as a CDK target by phospho-proteomics using a similar cyclin-Cdk fusion (Swaffer et al., 2016), cnp3 phosphorylation of the CDK consensus motif at S173 has been detected by mass spectrometry (Tay et al., 2019).

**Figure 4:**
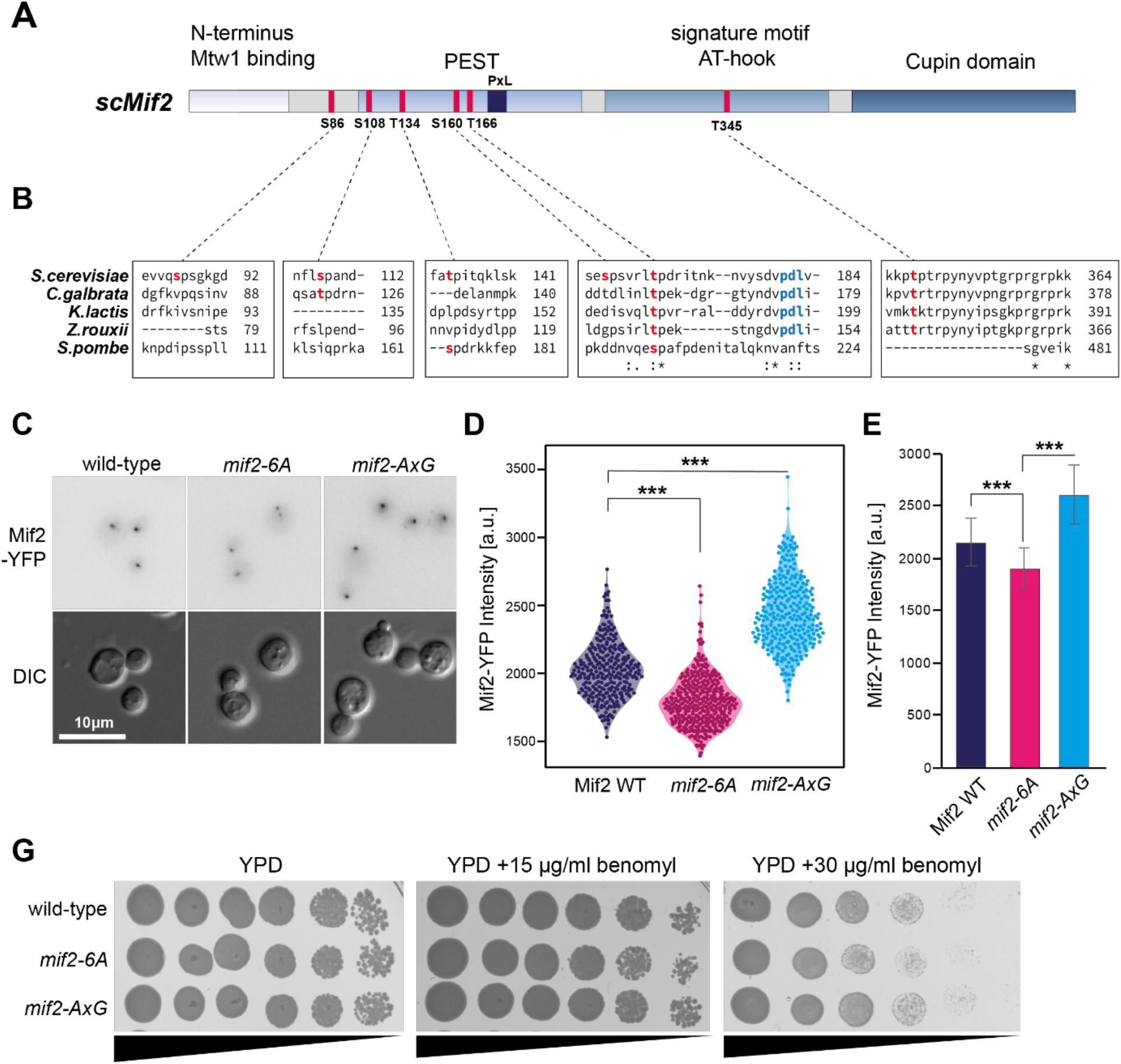
Mutation of putative CDK sites enriches Mif2 at kinetochores with minor effects on chromosome segregation. (A) Schematic of Mif2^CENP-C^ domain structure and putative CDK consensus sites. Consensus sites are highlighted in pink, a PxL-motif is shown in navy. (B) Sequence alignment of different yeast species shows that two CDK consensus sites within the PEST domain and the adjacent PxL motif are conserved across species. Alignment was performed using the Clustal Omega multiple sequence alignment tool. (C) Micrographs showing Mif2^CENP-C^ signal intensities in wild-type, *mif2-6A* and *mif2-AxG* cells. The scale bar is 10 µm. (D) Quantification of fluorescent signal intensities of Mif2^CENP-C^ wild-type, *mif2-6A* and *mif2- AxG* cells. Quantification was performed using FociQuant (Ledesma-Fernández, 2015). Statistical significance was assessed using ANOVA (*** = p-value < 0.005). (E) Quantification of fluorescence intensities shown as bar chart. Effects are subtle, but consistent. Error bars represent standard deviations, statistical significance was assessed using ANOVA (*** = p-value < 0.005). (G) Spot test analysis of wild-type, *mif2-6A* and *mif2-AxG* cells on rich media (YPD) or media containing the microtubule destabilising agent benomyl. *Mif2-6A* cells exhibit mild growth defects under high benomyl conditions. Black bars highlight cell dilution (wide end = high number of cells, narrow end = low number of cells). Each step accounts for a ten-fold dilution.

There is relatively little conservation between the amino acid sequences of human CENP-C and yeast Mif2, apart from the CENP-C signature motif. Consequently, these putative Mif2 CDK phospho-sites are not conserved in human CENP-C based on sequence alignment. However, we noted that CDK consensus sites are present in the functionally conserved PEST and DNA-binding domains of CENP-C, which were previously identified as targets of phospho- regulation (Nousiainen et al., 2006; Zhou et al., 2013).

To assess the importance of Mif2^CENP-C^ phosphorylation, we next generated phospho-mutant versions by replacing the six identified residues with alanine (*mif2-6A*) and, separately, an AxG mutant of the PxL motif, which may prevent early dephosphorylation by Cdc14 (Kataria et al., 2018). The mutant Mif2^CENP-C^ proteins were tagged with YFP (Figure 4C) and we quantified kinetochore fluorescence intensities of mutant cells compared to wildtype yeast. We found that *mif2-6A* kinetochore foci intensities were reduced in phospho-mutant yeast. In contrast, *mif2-AxG* yeast showed increased Mif2^CENP-C^ levels at kinetochores (Figure 4D-E and Supplementary file 6). Further, sensitivity to the microtubule-destabilising agent benomyl was mildly increased in *mif2-6A* cells compared to Mif2^CENP-C^ -WT and *mif2-AxG* yeast (Figure 4G). Notably, this effect was only visible at a high concentration of benomyl (30 µg/ml), indicating that CDK-mediated phosphorylation of Mif2^CENP-C^ alone is not crucial for chromosome segregation when microtubules are destabilised. This suggests that mutating some of the putative CDK sites is sufficient to impact Mif2^CENP-C^ kinetochore localisation. Although removal of the Cdc14 recognition site (PxL) increases Mif2^CENP-C^ levels, it does not reproduce the growth phenotype observed when recruiting sCDK (Figure 4).

Mif2^CENP-C^ is an essential, central component of the budding yeast kinetochore and structural studies have shown that Mif2^CENP-C^ spans the CCAN from the centromere and binds to the Mtw1^Mis12^ subunits of the outer kinetochore MIND complex (Figure 5A) (Hinshaw, 2019; Yan, 2019). Consequentially, we tested whether *mif2-6A* shows stronger growth defects in yeast cells with a compromised CCAN. We deleted different non-essential CCAN subunits in *mif2- 6A* cells and found that the deletion of the genes encoding the CTF19 complex subunits Iml3 and Ctf19 increased growth defects when cells were exposed to benomyl (Figure 5B). Notably, we were not able to generate viable *mif2-6A mcm21Δ* double mutant cells. Ctf19 is the kinetochore receptor for the cohesin-loader complex (Hinshaw, 2017), hence the increased benomyl sensitivity could be a combined effect of reduced cohesin and Mif2^CENP-C^ perturbation at kinetochores. We then measured spindle length in *ctf19Δ* and *mif2-6A ctf19Δ* cells using a Tub1-CFP tag (Figure 5C and Supplementary file 7) in mitotic spindles. There are typically two lengths of spindles seen in mitotic cells, first the more common metaphase spindles, which are typically <3 µm, and second, longer anaphase spindles. We applied a mixture model to describe spindle length as a bimodal distribution, with two components representing metaphase and anaphase spindles respectively (Scrucca et al., 2016). Mixture model fitting then allowed us to calculate average spindle lengths for both meta- and anaphase of the two genotypes. Interestingly, we found that spindle length was significantly increased in *mif2-6A ctf19Δ* cells in both, meta- and anaphase compared to *ctf19Δ* cells (Figure 5D). We next wanted to understand whether microtubule dynamics are affected in *mif2-6A ctf19Δ* cells. To this end, we followed spindle dynamics during mitosis using time-lapse microscopy (Figure 5E and Supplementary File 7) and observed that anaphase progression of *mif2-6A ctf19Δ* cells was significantly slower compared to *ctf19Δ* cells (Figure 5F).

**Figure 5:**
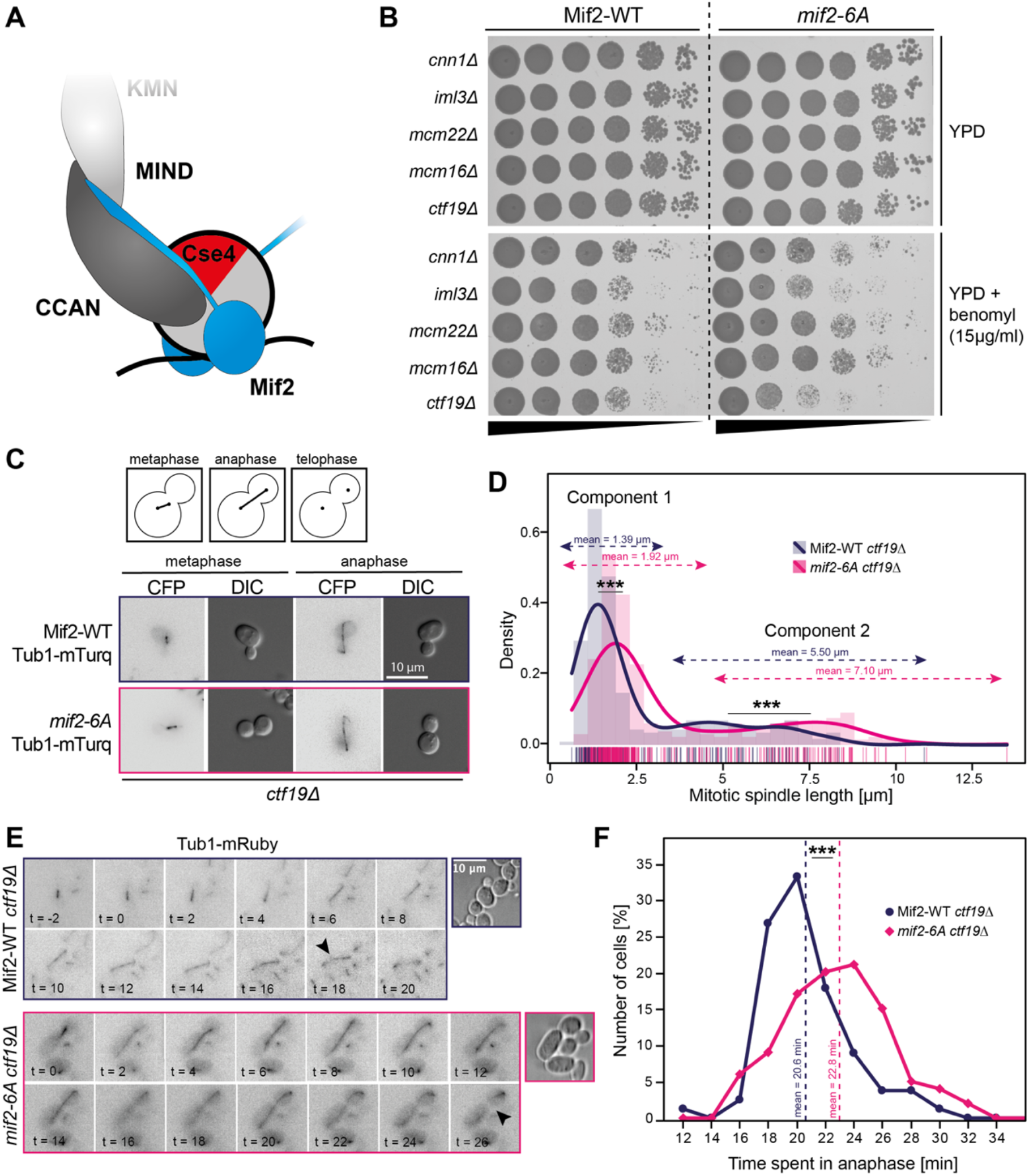
Mif2 phospho-mutants delay anaphase progression in a *CTF19* deletion strain. (A) Schematic of the budding yeast CCAN, highlighting Mif2^CENP-C^ in blue. A central domain of Mif2^CENP-C^ interacts with the centromeric nucleosome by binding to Cse4, whereas the N-terminus binds to the MIND component Mtw1^Mis12^ of the KMN network. Thus, Mif2^CENP-C^ spans the CCAN from the centromere to the outer kinetochore. (B) Spot test analysis of Mif2^CENP-C^ wild-type and *mif2-6A* yeast in various CCAN subunit deletion backgrounds (as stated on the left) with and without the microtubule destabilising agent benomyl. Increased benomyl sensitivity of *mif2-6A* cells was observed in IML3 and CTF19 deletion backgrounds. Spot test analysis was repeated 2 times to confirm phenotypes. Black bars highlight cell dilution (wide end = high number of cells, narrow end = low number of cells). Each step accounts for a ten-fold dilution. (C) Schematic and micrographs of spindle elongation from metaphase (short, bioriented) to anaphase (elongated) of *CTF19* deletion strains with wild-type or phospho-mutant Mif2^CENP-C^. The scale bar is 10 µm. (D) Analysis of spindle length of *CTF19* deletion strains with wild-type or phospho-mutant Mif2^CENP-C^. Bimodal spindle length distributions were dissected into two components using a mixture model to calculate the average spindle length specific for metaphase and anaphase cells. Mutating Mif2^CENP-C^ led to longer spindles in both metaphase and anaphase in *ctf19Δ* yeast. Single values are represented below the graph. Statistical comparison of means was performed using Student’s T-test (*** = p-value < 0.005). (E) Micrographs showing time-laps analysis of Mif2^CENP-C^ wild-type or mif2-6A in a *CTF19* deletion background. Black arrows highlight spindle disassembly. The scale bar is 10 µm. (F) Quantification of spindle elongation timing (from initial spindle extension to spindle disassembly). Spindle elongation time was extended in mif2-6A *ctf19Δ* cells. Statistical significance was assessed using Fisher’s Exact test (*** = p-value < 0.005).

In summary, our data suggest that CDK-mediated phospho-regulation of Mif2^CENP-C^ promotes Mif2^CENP-C^ localisation at kinetochores, which in turn supports kinetochore integrity. Our relatively mild phenotypes observed in combination with *CTF19* deletion underline how important spindle dynamics are throughout mitosis and indicate that multiple proteins and regulatory mechanisms work together to insure faithful chromosome segregation.

## Discussion

Major advances in identifying protein phosphorylation dynamics throughout the cell cycle have been made using phospho-proteomics (Godfrey et al., 2017; Holt, 2009; Swaffer, 2016; Touati et al., 2019; Ubersax et al., 2003). However, the role of individual protein phosphorylation events remains elusive in most cases. Low throughput phospho-deficient and -mimetic mutant analysis allows functional characterisation of protein phosphorylation but eliminates the dynamic aspect of post-translational modifications. In many cases, multiple sites or protein subunits of the same complex are targeted by cell-cycle phospho-regulators, hence mutation of residues within a single protein might not be sufficient to cause the requisite phenotypic changes in cell-cycle progression. In addition, different kinases and phosphatases often target the same proteins and recent studies have identified interplays of phospho-regulators at key targets such as the kinetochore (Klemm, 2021). In some cases, these kinases and phosphatases work in parallel to dynamically cycle phosphorylation status for highly regulated substrates (Saurin, 2018).

Alternative strategies to gain functional insights into specific phosphorylation events include the direct fusion of phospho-regulator subunits, such as cyclins, to potential substrate proteins, leading to constitutive phosphorylation (Kuilman et al., 2015), or using the SPI method to forcibly recruit phospho-regulators to cellular proteins (Mishra et al., 2021; Ólafsson & Thorpe, 2015, 2016, 2020). The advantage of these methods is that multiple sites and even surrounding proteins are targeted by the recruited or fused phospho-regulator. Here, we designed and generated a synthetic cyclin-dependent kinase that can be forced to any cellular protein tagged with GFP to promote targeted phosphorylation. We used the SPI method to recruit our synthetic kinase to proteins involved in chromosome segregation and identified a set of kinetochore, spindle pole and microtubule-associated proteins as sensitive to sCDK recruitment. Future research could express our synthetic kinase system in the *S. cerevisiae* GFP collection to unravel functional aspects of CDK phospho-regulation on a proteome-wide scale.

We focussed on one specific protein sensitive to sCDK recruitment independent of the spindle assembly checkpoint, the essential CCAN kinetochore component Mif2^CENP-C^. Although CENP-C phosphorylation by CDK promotes CENP-A binding in human cells (Watanabe, 2019), *S. cerevisiae* Mif2^CENP-C^ has not been identified as a CDK substrate. However, recent research has shown that Mif2^CENP-C^ is phosphorylated by Dbf2 and Cdc5 to promote inner kinetochore assembly (Hinshaw, 2023), kinases which often work in cooperation to drive mitotic progression (Klemm et al., 2021). Understanding the complexity and conservation of kinetochore phospho-regulation in a simple model organism allows easy characterisation of genetic interactions with other mutations, crucial to understand what drives cells from quiescent to dividing. We found that synthetic CDK recruitment leads to Mif2^CENP-C^ accumulation at kinetochores suggesting a similar role of CDK in budding yeast (Figure 3). Interestingly, CDK recruitment drives rapid progression into anaphase after release from metaphase arrest, but cells ultimately fail to divide and arrest in a large-budded state, indicating that centromeric dephosphorylation during mitotic exit may be crucial for completing cell division. However, due to the nature of the SPI system, neighbouring proteins in vicinity of Mif2^CENP-C^ could also be phosphorylated by Mif2^CENP-C^ associated sCDK, promoting a phenotype not exclusively produced by Mif2^CENP-C^ phosphorylation.

Mutating putative CDK sites of Mif2^CENP-C^ reduces Mif2^CENP-C^ kinetochore levels (Figure 4) indicating less efficient kinetochore recruitment, Cse4 binding, or protein stability. Previous studies have estimated that two to four copies of Mif2^CENP-C^ are present at each yeast kinetochore and that Mif2^CENP-C^ dimerisation is crucial for proper kinetochore function (Cohen, 2008). As *mif2-6A* mutant cells grow efficiently and are largely tolerant to microtubule destabilisation, we assume that mutating CDK sites does not affect Mif2^CENP-C^ dimerization. Rather, we speculate that the decrease in fluorescence intensity observed in *mif2-6A* mutants is based on a reduced pool of unbound Mif2^CENP-C^ subunits at kinetochores, which could be targeted for degradation based on their phosphorylation status. Similarly, a recent study also reported a crucial role of DDK and Cdc5 for Mif2^CENP-C^ Cse4 interaction by phosphorylation sites within the PEST domain of Mif2^CENP-C^ (Hinshaw, 2021). As most of the putative CDK sites are also present within the PEST domain we suggest an interplay between CDK, DDK and Cdc5 at Mif2^CENP-C^ to regulate Mif2^CENP-C^ recruitment and binding to Cse4 at the budding yeast kinetochore. We propose that Mif2^CENP-C^ phosphorylation prevents ubiquitination and subsequent degradation when Mif2^CENP-C^ is stably incorporated into kinetochores. Unbound Mif2^CENP-C^ would then be rapidly dephosphorylated by Cdc14 recruitment via the PxL domain, hence mutating PxL stabilises Mif2^CENP-C^ and increases Mif2^CENP-C^ foci intensities at kinetochores. This hypothesis is supported by previous studies which reported crosstalk between phosphorylation and ubiquitin-mediated degradation (Hunter, 2007). Notably, a recent study identified CDK-mediated phospho-degrons within subunits of the CTF19 complex, indicating a tight relationship between phospho-regulation and protein stability (Böhm, 2021).

We also observed a genetic interaction between *MIF2* and *CTF19*, as *mif2-6A ctf19Δ* cells show increased benomyl sensitivity and an altered spindle phenotype (Figure 5). It remains unclear if this phenotype is based on the role of Ctf19 as the kinetochore receptor for the cohesin loader complex, or whether Mif2^CENP-C^ phosphorylation directly affects centromeric cohesin. Interestingly, *mif2-6A ctf19Δ* cells are enriched for longer anaphase spindles and delayed spindle dissociation it will be interesting to determine whether *mif2-6A* affects spindle disassembly. Our data suggest an important role of CDK at the budding yeast CCAN and, together with the recent findings of Hinshaw and colleagues imply a dynamic and complex model of Mif2^CENP-C^ phosphorylation during cell cycle progression.

Our synthetic kinase and in-silico analysis of phosphorylation sites strongly suggest that Mif2^CENP-C^ is a direct target of CDK phospho-regulation and that CDK-mediated phospho- regulation of Mif2^CENP-C^ promotes Mif2^CENP-C^ localisation at kinetochores, which in turn supports kinetochore integrity. Our system proved to be a sensitive methodology to identify new targets of cell cycle phospho-regulation which opens the way for broader use of this approach to study cell cycle control in the future.

## Material and methods

### Yeast Methods

All yeast strains used in this study are listed in Table S1 and plasmids are listed in Table S2 (Supplementary file 8). Green fluorescent protein (GFP) and yellow fluorescent protein (YFP) strains are based upon BY4741 (*his3Δ1 leu2Δ0 met15Δ0 ura3Δ0)* (Brachmann et al., 1998; Huh et al., 2003). The universal donor strain (UDS) is a derivative of W303 (*can1-100 his3-11,15 leu2-3,112 ura3-1 RAD5*) (Zou & Rothstein, 1997). Yeast cells were cultured in standard growth or auxotrophic selection medium with 2% carbon source (Sherman, 2002). SPI plasmids were generated by gap-repair cloning directly in yeast or by using the NEBuilder HiFi assembly kit (New England Biolabs) by combining linearised plasmid with gene fragments with homologous ends. Mutant gene fragments were synthesised by GeneArt (ThermoFisher) and transformed into BY4741 yeast. All plasmids and strains were validated by Sanger sequencing (GENEWIZ Brooks Life Science, UK).

### Synthetic Physical Interactions

SPI screens were performed as described in previous studies (Ólafsson, 2015; Ólafsson & Thorpe, 2018). Arrays of GFP strains were transformed separately with either control or experimental plasmids. Selective ploidy ablation (SPA) was used to introduce plasmids into arrays of yeast strains consisting of 89 members of the GFP collection that represent proteins that are expressed during mitosis (Huh, 2003; Reid et al., 2011). Briefly, the SPA method utilises a Universal Donor Strain (UDS, W8164-2B), which contains conditionally active centromeres with a *URA3* cassette and *GAL1* promoter adjacent to each centromere and is transformed with each of the SPI plasmids. These donor strains were then mated with members of the GFP collection on rectangular agar plates using a pinning robot (ROTOR robot, Singer Instruments, UK) in 4 replicates and 1536 colonies per plate. The resulting diploids were then sequentially selected to maintain the query strain GFP genome and plasmid while destabilising and then removing the chromosomes of the UDS by growing the cells in medium containing both 5-Fluoroorotic acid and galactose. Finally, the plates were scanned using a desktop flatbed scanner (Epson V750 Pro, Seiko Epson Corporation, Japan).

### SPI Data analysis

Colony sizes on SPI screening plates were measured using the colony measurement engine tool for ImageJ (Dittmar et al., 2010) and the resulting data was analysed using the open-source software ScreenGarden (Klemm et al., 2022), which calculates mean log growth ratios (LGRs) of experimental and control plates for each interaction. LGRs are calculated as the natural logarithm of control/experiment colony size for each colony and each control individually and then averaged over the replicates and controls. GFP strains with an average colony size of less than 30% compared to the plate median after forced association of the GBP control were defined as generally sensitive to protein recruitment and were excluded from further analysis. High-confidence SPIs were further defined based on ScreenGarden’s q-value evaluation of replicates (Klemm et al., 2022).

### Microscopy

We used epifluorescence microscopy to determine the cellular localisation of FP-tagged proteins and colocalisation. Cells were grown overnight, diluted 1/5, grown for 2 hours in selection or synthetic complete media with 2% w/v glucose, and either imaged immediately or treated as described. Strains containing SPI constructs under the control of the *MET3* promoter (pMET3) were grown with excess methionine (>2 mM) to prevent plasmid expression and pMET3 expression was induced in 10 µM methionine. Before imaging, cells were mixed with 0.7% low melting point agarose in growth medium on glass microscope slides. A Zeiss Axioimager Z2 microscope (Carl Zeiss AG, Germany) was used to image cells using a 63x 1.4NA apochromatic oil immersion lens. Fluorescence was excited using a Zeiss Colibri LED illumination system (GFP=470 nm, YFP=505 nm, and RFP=590 nm) and differential interference contrast (DIC) prisms were used to enhance the contrast in bright field. The emitted light was captured using a Hamamatsu Flash 4.0 Lte camera with an FL-400 CMOS sensor (6.5 µm pixels, binned 2x2). Exposure times were adjusted to ensure that signal intensities remained below saturation and remained identical between control and experimental images unless stated otherwise. Images were acquired using the Zen software (Zeiss) and analysed and prepared using the Icy BioImage Analysis unit (version 2.0.3.0) [87] and FIJI/ImageJ (Schindelin et al., 2012).

For time-lapse microscopy, cells were grown to log-phase, transferred onto agar pads and moved to imaging chambers. After 1 hour of incubation at 30°C, cells were imaged using a DeltaVision Elite microscope (GE Healthcare, USA) with a 60× 1.42NA Oil Plan APO and an InsightSSI 7 Colour Combined Unit illumination system (mRuby2 = 575 nm). Images were captured with a front-illuminated sCMOS camera, 2,560 × 2,160 pixels, 6.5 *μ*m pixels, binned 2×2. Time-lapse videos were captured over 2 hours, with images captured at 2-minute intervals. Images were analysed using FIJI/ImageJ (Schindelin, 2012).

### Western blotting

3xFLAG-tagged Mif2^CENP-C^ cells were harvested after 4 hours of growth at restrictive (37°C) or permissive (23°C) temperature and whole cell lysates were prepared using glass beads and a cell disrupter (Scientific Industries Inc.) in Laemmeli buffer containing an EDTA-free protease inhibitor cocktail (Thermo Fisher). Samples were subsequently denatured at 95°C and loaded onto 7.5% SDS-gels with 50 µmol/ml phostag agarose to separate proteins based on their charge. A Phostag-free gel was run analogously as a loading control. Western blotting was used to transfer protein into PDFV membranes. After antibody incubation (Anti-FLAG^®^ M2, Sigma Aldrich), bands were visualised using ECL select (Cytiva) or Clarity Western ECL substrate (BioRad) in a ChemiDoc gel imaging system (BioRad).

### Statistical analysis

All statistical methods shown in graphs are described in the respective figure legends. SPI and other averaged data are shown as mean over 16 replicates with standard deviation unless stated otherwise. For continuous data, statistical significance was assessed using a two-tailed unpaired Students T-test or ANOVA, in case of multiple comparisons. For discrete data, Fisher’s Exact Test was used to test significance. P-values are represented as follows: * = p- value < 0.05, ** = p-value < 0.01, *** = p-value < 0.005.

### Online supplemental material

**Figure S1** shows the colocalization of SPI constructs to the spindle body protein Spc42. **Figure S2** shows shows the activity and recruitment of active and inactive sCDK SPI constructs. **Figure S3** shows the correlation between sCDK recruitment and Mif2^CENP-C^ kinetochore localisation and sCDK-mediated occurrence of additional kinetochores. **Figure S4** shows altered Mif2^CENP-C^ phosphorylation and Mif2^CENP-C^ intensity at kinetochores in yeast expressing temperate- sensitive alleles of Cdc28 and Cdc14. **Supplementary file 1** includes mean LGRs and colony sizes of CDK SPI screens with components involved in chromosome segregation. **Supplementary file** 2 contains SPI data from mad3 deletion SPI screens. **Supplementary file** 3 contains Mif2^CENP-C^ intensity data with and without sCDK recruitment. **Supplementary file 4** reports the number of large-budded cells and cell cycle stage after release from nocodazole with and without sCDK recruitment. **Supplementary file 5** contains the quantification of bands from phostag analysis and Mif2^CENP-C^ intensities at restrictive and permissive temperatures of cells expressing temperature-sensitive alleles of Cdc28 or Cdc14. **Supplementary file 6** reports kinetochore intensities of Mif2^CENP-C^ and phospho-regulation mutants. **Supplementary file 7** contains the data showing that mutation of putative CDK sites within Mif2^CENP-C^ to alanine leads to increased spindle length and anaphase progression in *CTF19* deletion backgrounds. **Supplementary file 8** lists all strains and plasmids used in this study.

### Data availability

All data used to conclude our findings are found in the article and/or supplemental material. Raw SPI data outputs not listed in supplementary files for clarity are available upon request from the corresponding author. R and Perl scripts for data analysis are reposited on the author’s GitHub (https://github.com/CinziaK/).

## Acknowledgements

We thank Frank Uhlmann and the late Angelika Amon for kindly providing the *Δclb1, Δclb3, Δclb4, clb2-ts* yeast strains to test activity of the sCDK system. We also thank Frank Uhlmann for insightful discussions during different stages of the project. Further, we would like to thank all past and present members of the Thorpe lab for helpful conversations and support and Damien Coudreuse for comments on this manuscript.

We would like to thank all our funding bodies, Queen Mary University (CKs studentship and consumables), the Francis Crick Institute, which receives its core funding from Cancer Research UK (FC001183), the UK Medical Research Council (FC001183), and the Wellcome Trust (FC001183); also the BBSRC (BB/R00868X/1 & BB/T017716/1). These funding bodies played no roles in the design of the study and collection, analysis and interpretation of data and writing the manuscript. The authors declare no conflict of interest.

## Author contributions

P. H. Thorpe designed the research project, evaluated results and supported writing and revising the manuscript. C. Klemm and G. Olafsson designed and performed experiments. C. Klemm performed statistical data analysis and wrote the manuscript. All authors have read and approved the manuscript.

## Corresponding author

Correspondence to Peter H. Thorpe.

## Supplement figures

**Figure S1:**
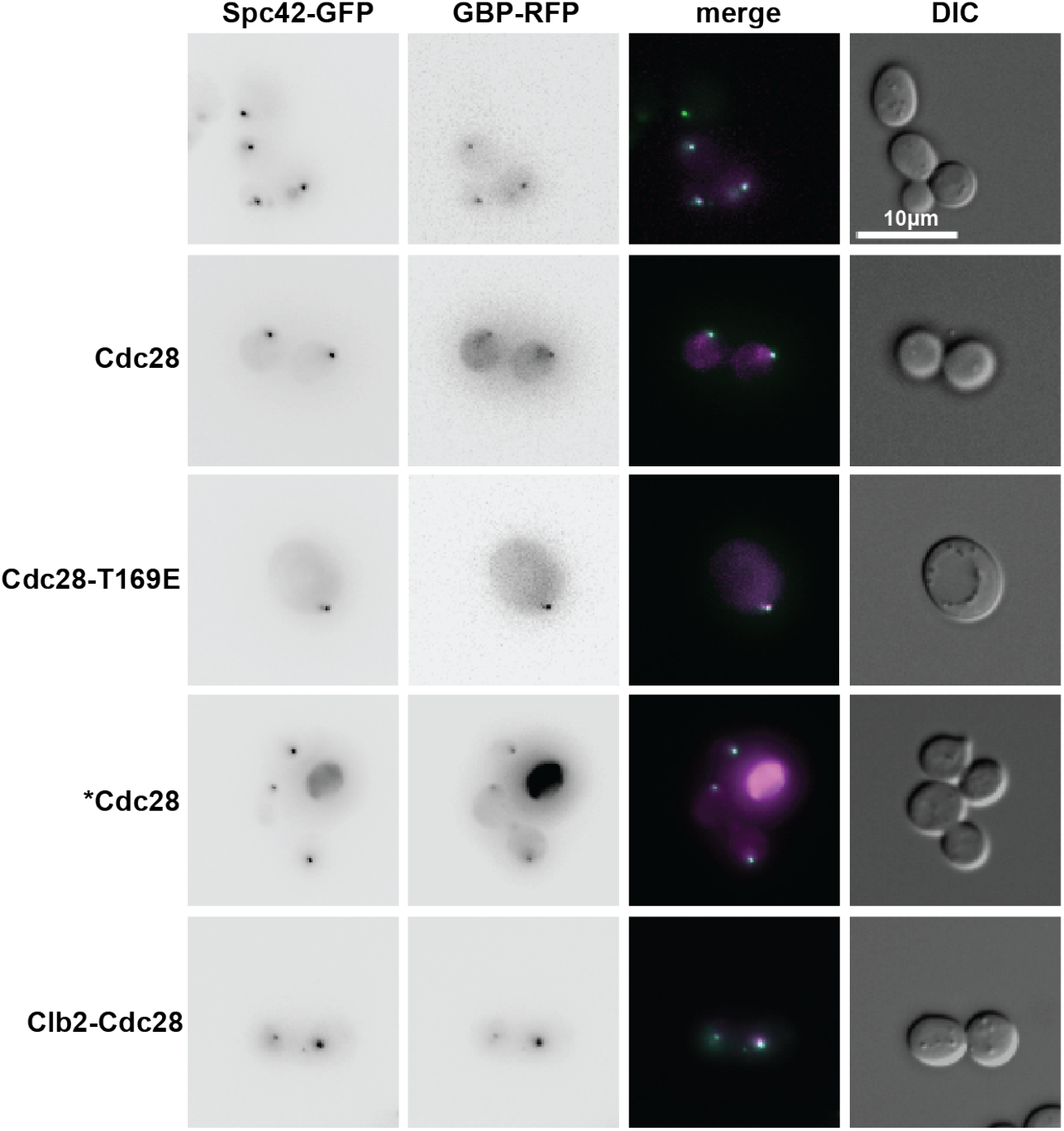
Colocalisation of CDK SPI constructs. Micrographs showing the colocalisation of Cdc28, Cdc28-T169E, *Cdc28 and Clb2-Cdc28 and the spindle pole body component Spc42. Spc42 was chosen as representation due to its distinct cellular localisation and easy detection. The scale bar is 10 µm.

**Figure S2:**
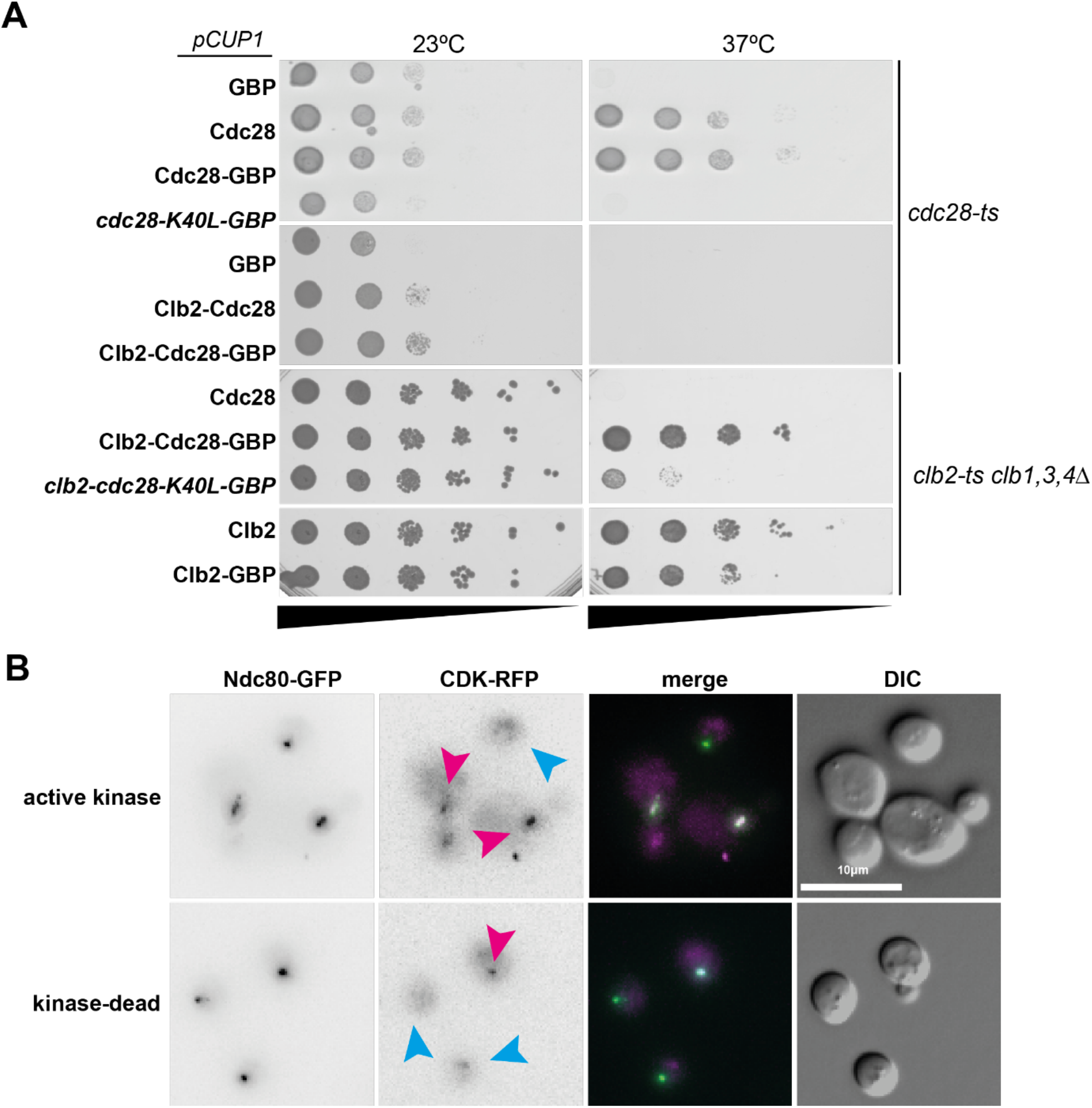
Clb2-Cdc28 is insufficient in driving the cell cycle and undetectable in G1. (A) Serial dilution spot assay showing that Clb2-Cdc28 and Clb2-Cdc28-GBP cannot rescue cdc28-ts lethality at restrictive temperature. Cdc28 and Cdc28-GBP function as positive controls and GBP recruitment is used as negative control. This is likely a result of insufficient binding of Clb2-bound Cdc28 to other cyclins and/or rapid Clb2-Cdc28 degradation. (B) Micrographs showing successful forced protein recruitment of active and kinase-dead Clb2-Cdc28-GBP fusions to the kinetochore protein Ndc80. Notably, colocalisation was only observed in budded cells (indicated by pink arrows) and not in G1 cells (blue arrows). The scale bar is 10µm.

**Figure S3:**
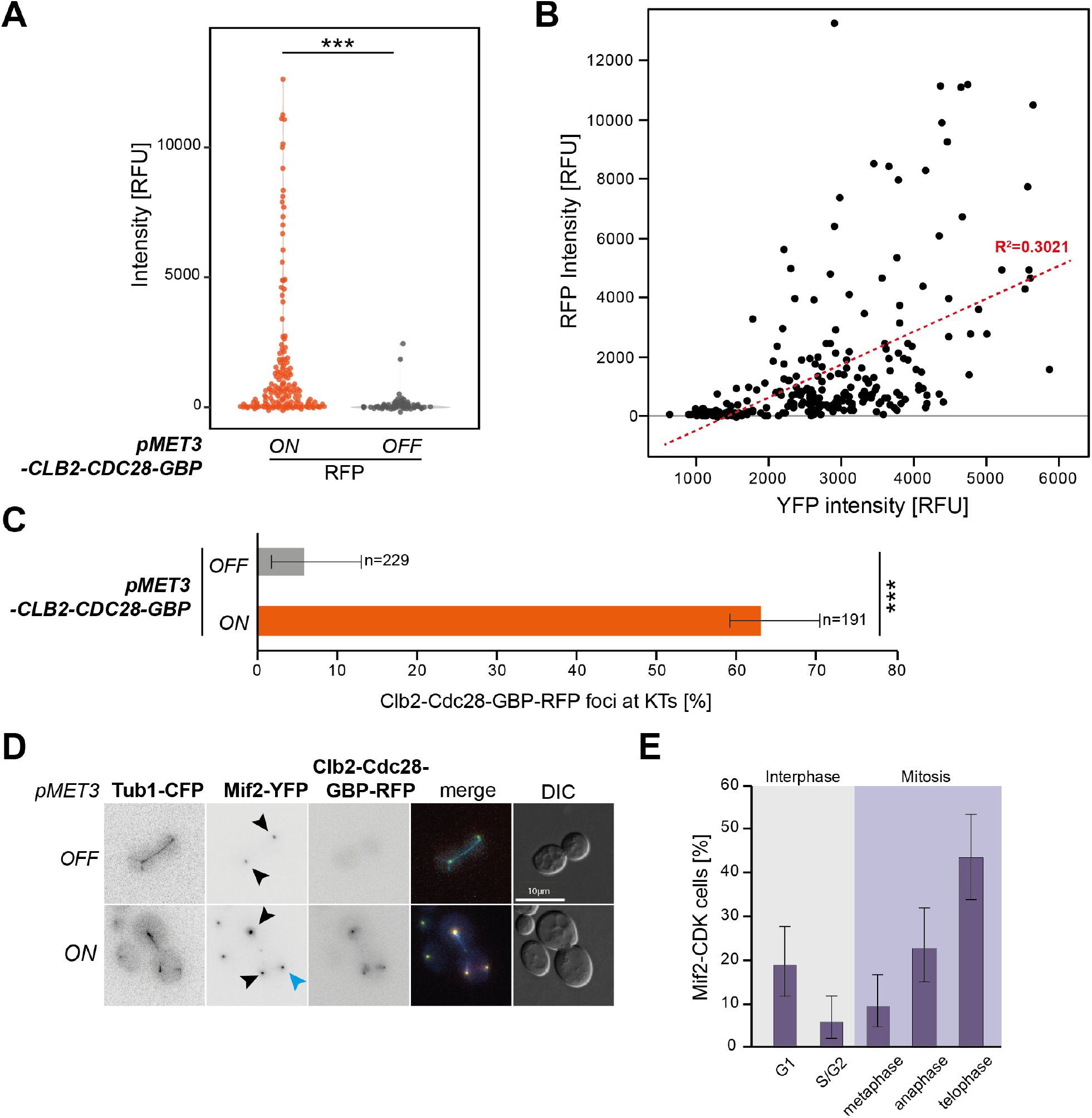
Fluorescence intensities of Clb2-Cdc28-GBP-RFP and Mif2-YFP are correlated. (A) The violin plots show Clb2-Cdc28-GBP-RFP signals when *pMET3-CLB2-CDC28* expression is induced (10µg/ml methionine) or repressed (2mM methionine). Signal intensities vary in *pMET3 ON* conditions and RFP signals are not always detectable. Quantification of fluorescence intensities using the semi-automated FociQuant image analysis software (Ledesma-Fernández & Thorpe, 2015). Statistical significance was assessed using a Students T-test (*** = p-value < 0.005). (B) The correlation between Mif2-YFP and Clb2-Cdc28-GBP-RFP signals is shown in the scatterplot. Cells with strong RFP signals also show increased YFP signals. A linear regression is shown with the red dotted line, the grey line is the zero line for RFP signal intensity. R^2^ is the coefficient of determination and indicates the correlation between RFP and YFP signal intensities. (C) Percentage of Mif2-YFP foci with detectable levels of Clb2-Cdc28-GBP-RFP. Approximately 5% of cells showed RFP co-localisation to YFP signals when *pMET3* expression is repressed (*OFF*, 2mM methionine), whereas *pMET3* induction (*ON*, 10µg/ml methionine) resulted in 63% of cells with detectable RFP signals. RFP detection threshold were set to an intensity of 200. Error bars are calculated as 95-percentiles. Statistical significance was assessed using a Fisher’s Exact test (*** = p-value < 0.005). (D) An additional copy of the tubulin Tub1 tagged with cyan fluorescent protein (CFP) shows mitotic spindle integrity in Mif2-YFP cells with and without CDK-RFP recruitment. Notably, some cells presented extra kinetochore foci. The scale bar is 10 µm. (E) Quantification of cell cycle stage based on cell shape (budded, unbudded) Mif2-YFP and Tub1-CFP signal shows that cells are enriched in telophase, with disassembled spindles but prior to cytokinesis, after forced CDK association.

**Figure S4:**
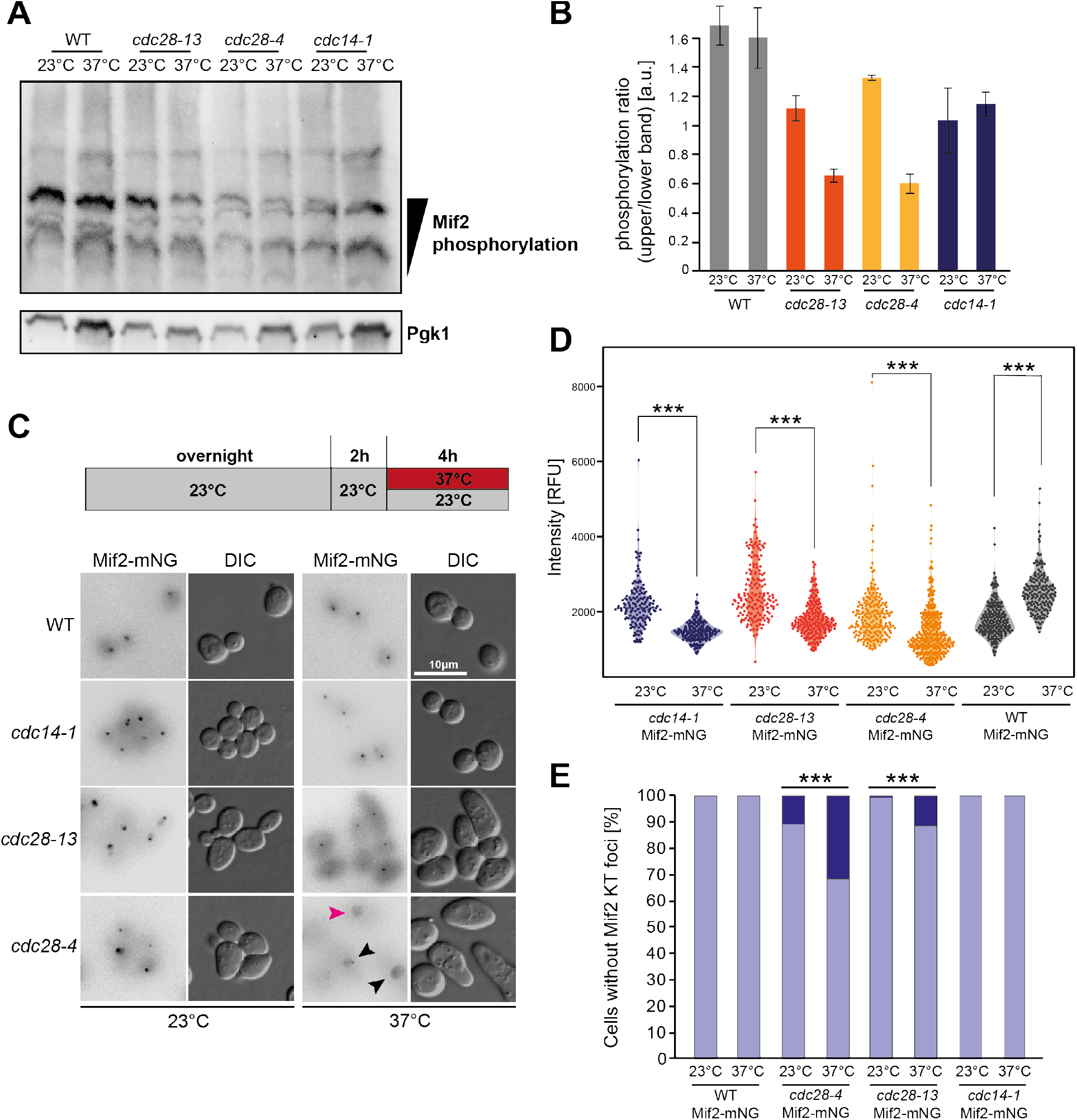
Depletion of Cdc28 function leads to decrease in Mif2 phosphorylation and recruitment to kinetochores. (A) Western blot of phostag-agarose SDS-page, separating Mif2^CENP-C^ molecules based on their phosphorylation status. The slower-migrating bands represent increased phosphorylation, whereas faster-migrating bands account for less phosphorylation. (B) Quantification of western blot analysis shows decrease in phosphorylation ratio (upper/lower band) of Mif2^CENP-C^ at restrictive temperature in cdc28-ts strains. Error bars represent standard deviation between replicates. Statistical significance was assessed using Student’s t-test. (C) Micrographs showing Mif2^CENP-C^ signals in temperature sensitive alleles of Cdc28 or Cdc14 at permissive (23°C) or restrictive (37°C) temperature. The scale bar is 10 µm. (D) Quantification of kinetochore fluorescence intensities using FociQuant (Ledesma-Fernández, 2015). Statistical significance was assessed using ANOVA (*** = p-value < 0.005). (E) Quantification of cells with kinetochore signals. Error bars represent 95% confidence intervals, statistical significance was assessed using Fisher’s Exact test (*** = p-value < 0.005).

## Notes

### Competing Interest Statement

The authors have declared no competing interest.

